# Optimal Point Process Filtering and Estimation of the Coalescent Process

**DOI:** 10.1101/024737

**Authors:** Kris V Parag, Oliver G Pybus

## Abstract

The coalescent process is an important and widely used model for inferring the dynamics of biological populations from samples of genetic diversity. Coalescent analysis typically involves applying statistical methods to either samples of genetic sequences or an estimated genealogy in order to estimate the demographic history of the population from which the samples originated. Several parametric and non-parametric estimation techniques, employing diverse methods, such as Gaussian processes and Monte Carlo particle filtering, already exist. However, these techniques often trade estimation accuracy and sophistication for methodological flexibility and ease of use. Thus, there is room for new coalescent estimation techniques that can be easily implemented for a range of inference problems while still maintaining some sense of statistical optimality.

Here we introduce the Bayesian Snyder filter as a natural, easily implementable and flexible minimum mean square error estimator for parametric demographic functions. By reinterpreting the coalescent as a self-correcting inhomogeneous Poisson process, we show that the Snyder filter can be applied to both isochronous (sampled at one time point) and heterochronous (serially sampled) estimation problems. We test the estimation performance of the filter on both standard, simulated demographic models and on a well-studied empirical dataset comprising hepatitis= C virus sequences from Egypt. Additionally, we provide some analytical insight into the relationship between the Snyder filter and popular maximum likelihood and skyline plot techniques for coalescent inference. The Snyder filter is an exact and direct Bayesian estimation method that provides optimal mean square error estimates. It has the potential to become as a useful, alternative technique for coalescent inference.

## 1. Introduction

Genetic sequences contain information about the dynamics of the population from which they were sampled. The coalescent process provides a framework for extracting this information by describing the shared ancestry among *n* individuals randomly sampled from a population of effective size *N*(*t*) **≫** *n* [1]. The shared ancestry of the sampled individuals can be visualised as a random, ultrametric, bifurcating genealogy with *n* tips and *n –* 1 branches. The branch lengths give the times at which sampled lineages coalesce. These coalescence times depend on *N*(*t*) which is also called the demographic function and describes the dynamics of the population. A key problem in coalescent inference is the estimation of *N*(*t*) or its parameters from an empirically observed genealogy, or a set of sampled genetic sequences.

The original, standard coalescent was developed by Kingman for a constant *N*(*t*) and for sets of genetic sequences that are sampled at one time point (isochronous sampling) [1]. Since then, the coalescent model has been generalised to incorporate deterministically varying population sizes [2], stochastic population fluctuations [3], geographically structured populations [4], and data sets containing sequences sampled at different time points (heterochronous sampling) [5]. As a result, the coalescent model has been applied to a range of problems in many biological disciplines including conservation biology, anthropology and epidemiology [6]. Our work is geared towards infectious disease epidemiology where pathogen populations, due to their large size and very rapid molecular evolution, are often deterministically varying in size and heterochronously sampled. In this setting, the coalescent process has been successfully used to infer the growth and history of the hepatitis C epidemic in Egypt [7], the oscillating behaviour of dengue virus in Vietnam [8] and to estimate the generation time of HIV-1 within individual infected patients [5]. The accuracy and efficiency of such inferences are linked to the statistical techniques used. Consequently, the design of good coalescent demographic inference methods is important [9].

We focus on the coalescent inference problem for a haploid population with deterministically varying population size, under both isochronous and heterochronous sampling. We follow the standard coalescent assumptions of a panmitic (well mixed) and neutrally evolving population that is sparsely and randomly sampled [10]. Several methods for inferring the demographic function, *N*(*t*), have been developed and can be broadly categorised into parametric and non-parametric approaches. The former characterises *N*(*t*) using a biologically-inspired function with a fixed number of demographic parameters. These parameters interact in a predefined manner and the dimensionality of the model is independent of *n*. In contrast, non-parametric methods use less constrained functions for *N*(*t*) and therefore make weaker assumptions about demographic dynamics. This allows a more robust and generalised description of population size. However, this comes at the expense of less statistical power, and with the possibility of model dimensionality increasing with *n* [11].

Among non-parametric coalescent inference techniques, the skyline plot [12] and its variants, such as the skyride [13] and skygrid [14] plots have become popular [6]. The original version, now known as the classic skyline plot [12], computes harmonic mean estimates of *N*(*t*). However, this approach is very noisy because it uses a single coalescent event to generate a piecewise constant *N*(*t*) estimate for each inter-node interval (the period between successive coalescent events in a genealogy). Generalisations of the skyline plot pool the periods between coalescent events in order to reduce noise. These methods utilise change point models [15], Gaussian Markov random fields [13] or Akaike information measures [6] but still retain a mean-type estimator for *N*(*t*). Other more recent non-parametric techniques use machine learning principles, such as Gaussian processes [16], to describe inter-node covariation. These fit data to mean and covariance functionals, and so make use of more of the demographic information which is embedded within the coalescent process.

While these newer non-parametric approaches have increased statistical power and reduced noise, they tend to be methodologically and computationally complex [13]. Further, these more sophisticated extensions can be seen as implicitly more parametric in formulation, since they impose more constraints on the estimator space and require more information. Consequently, in this study, we focus on parametric estimation. Specifically, we assume that a suitable demographic model 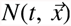 has already been chosen and that its parameters 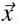, or a function of them, are to be estimated from the data in an optimal way. In this paper, we introduce and analyse the Snyder filter [17], a technique from electrical and systems engineering, as a means of solving this estimation problem. The Snyder filter is an explicit, parametric, Bayesian inference technique that directly solves dynamical equations for the joint posterior distribution of 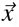. These equations can then be used to obtain a conditional mean estimator which minimises the mean square error between the true parameter (or function) and its estimate.

The Snyder filter is unlike other existing Bayesian methods for coalescent inference, which typically approximate the posterior distribution by using Markov Chain Monte Carlo (MCMC) or importance sampling [18] [19]. These approaches, while able to account for genealogical uncertainty, can be complex or difficult to implement [20], especially when they involve integration over complicated high dimensional parameter spaces. Notably, because the Snyder filter directly computes the posterior distribution, is it not affected by problems related to MCMC convergence, which have been particularly acute for some coalescent models [21]. We show how the Snyder filter, which treats coalescent data as a point process stream, can be used as an alternative and useful Bayesian estimator. We also clarify the relationship between Snyder filtering and standard maximum likelihood (ML) analysis as well as explore its connection to the classic skyline plot. The Snyder filter has remained largely unknown to the biological sciences and, to our knowledge, has only been applied to neuronal spiking by Bobrowski *et al* [22] and to invertebrate visual phototransduction by Parag [23]. At present, our framework does not yet account for genealogical uncertainty. As a result, we focus on the problem of estimating demographic parameters from a single genealogy.

We first show how the coalescent process, with time varying population size, can be reinterpreted as a self-correcting, inhomogeneous Poisson process and then develop the appropriate Snyder filter for both heterochronous and isochronous sampling. Next, we derive an analytic Snyder estimator for the standard Kingman coalescent, which helps clarify the relationship between this Bayesian technique and standard ML approaches. We also develop the link between Snyder filters and classic skyline plots in an Appendix. This completes our theoretical treatment of the filter. We then explore and quantify the performance of the Snyder filter by applying it to (i) data simulated under several canonical, deterministically time-varying, demographic models and (ii) a well-studied empirical dataset comprising hepatitis C virus (HCV) gene sequences from Egypt. This HCV dataset has been extensively used in previous studies and thus allows us to compare our method with existing approaches.

## 2. Methods

### 2.1. The Coalescent as a Self-Correcting Poisson Process

Let *n* ≥ 2 samples be taken from a population of size *N(t)* ≫ *n* at *K* distinct times so that 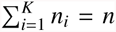. The pair (*s*_*i*_, *n*_*i*_) represents the i^th^ sample time and the corresponding number of tree tips (sampled lineages) introduced at that time. Here *s*_1_ = 0 is always taken as the present (the most recent time of sampling). Time therefore increases into the past. If *K* = 1, then sampling is isochronous. The coalescent process for *n* lineages is usually described as an inhomogeneous Poisson process with a maximum count of *n –* 1. In this interpretation, the stream of *n* – 1 coalescent events is separated into intervals that end with either a coalescent or sampling event. The *k*^th^ coalescent time, *c*_*k*_, is the time point at which the lineage transition *k → k –* 1 occurs. The final coalescent event is therefore at *c*_2_ and the first at *c*_*n*_ > 0. The coalescent rate is described for each interval by conditioning on the number of lineages in that interval [2]. All the information for estimation is encoded in the coalescent event times, *(c*_*k*_, *k*) for 2 ≤ *k ≤ n* and in the knowledge of the sampling scheme, (*s*_*i*_, *n*_*i*_) for 1 ≤ *i ≤ K*. In this work we focus on estimation given the values of *c*_*k*_, *s*_*i*_ and *n*_*i*_ and we do not consider the bifurcating genealogy from which they arise.

We extend this standard description by embedding both the fall in lineages (due to stochastic coalescent events), and the discontinuous rise in lineages (due to sampling), explicitly within the Poisson rate. This removes the need for direct conditioning and allows the coalescent to be redefined as a self-correcting inhomogeneous Poisson process between sampling times. Hence changes in the number of lineages are treated as Poisson rate feedback. In self-correcting processes previous events inhibit the occurrence of following events. The coalescent process exhibits this behaviour because the rate of producing new events falls with the event count as time increases into the past. This self-correcting property is independent of the form of *N*(*t*) since the standard Kingman coalescent can be recovered from the variable population size case by appropriate time rescaling [2]. We define *u*(*t*) as the number of coalescent events from the present (time 0) to time *t* in the past, and *h*(*t*) as the count of the total number of samples introduced into the process over that time period (see equations 1 and 2). Then max_*t*_ *h*(*t*) = *n = h(t* ≥ *s*_*K*_) (the number of samples or tree tips) and maxt *u*(*t*) = *n –* 1 = *u*(*t ≥ c*_2_) (the total number of coalescent events). For isochronous sampling *h*(*t*) = *n*_1_ = *n* at all *t*. The indicator function 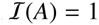 if condition *A* is true and is 0 otherwise.

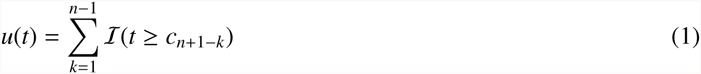

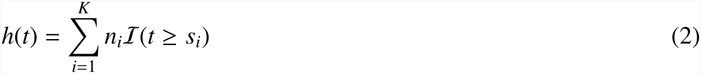

The difference *r*(*t*) = *h*(*t*) – *u*(*t*) is the number of extant lineages at time *t*. If 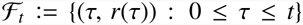 is the counting process with points at both sampling and coalescent event times, then it contains all the causally observable information from 0 to *t*. The feedback dependent (self-correcting) coalescent rate at time *t* is written 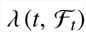 (or simply ***λ***) and is defined as:

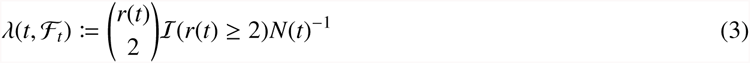

The coalescent inference problem is therefore equivalent to estimating the rate parameters of a self-correcting inhomogeneous Poisson process, which is also a doubly stochastic Poisson process (because it has a stochastic intensity). This key insight allows us to apply the filtering techniques of Snyder [24].

### 2.2. Optimal Snyder Filtering

The Snyder filter is an exact Bayesian filter and makes no approximations on either the observed or estimated process. It optimally reconstructs the informed posterior of the parameters 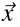 of the underlying Poisson process intensity, *λ*, given observations 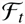 and priors on 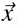 [24]. It was originally developed for doubly stochastic Poisson processes and continuously solves ordinary differential equations for the model posteriors between observed event times. It implements discontinuous posterior updates at event times. Since the coalescent process fits this framework, the Snyder filter can be adapted to estimate the demographic function and its parameters. As input, the filter only requires the observed coalescent times, *c*_*k*_, the sampling scheme, (*s*_*i*_, *n*_*i*_), and priors on the demographic parameters. We assume these are all known without error. The filter outputs posterior distributions from which conditional mean estimates can be calculated. Its computational complexity depends on the number of parameters to be estimated, l, the number of tree tips, *n* and the dimension of the filter, *m* (which is defined below). As the filter only requires the integration of linear, ordinary differential equations it is easy to implement and avoids the non-linear optimisation or MCMC marginalisation which can make other coalescent inference methods unwieldy.

The Snyder filter is defined as follows. Let a general multivariate demographic function with *l* parameters of true values, 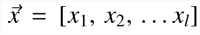, be denoted as 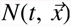. Parameter *x*_*i*_ takes a value somewhere in the domain *χ*_*i*_. This domain is gridded into *m*_*i*_ discretised points, and it is only this space and a prior on it that are available to the inference method. The joint prior, 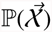 is defined on a space of 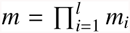 points with 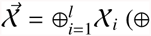 indicates a Cartesian product). The filter solves a differential equation on each point and so has dimension *m*. The posterior vector, 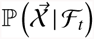 has *m* components and is Cartesian ordered on the space of 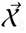. Let the unnormalised version of this vector and its *j*^th^ component be *q*\*(*t*) and 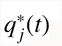. The normalised forms are then *q*(*t*) and 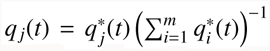 respectively. The use of unnormalised probabilities, as developed in [25], converts the original non-linear Snyder differential equations into a linear set that, once solved, need only be scaled to 1. If 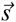 is the vector of *K* sample times and 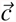 the vector of *n –* 1 coalescent times and an additional time *c*_*n+*1_ *= s*_1_ = 0, then the total number of distinct non-zero event times is *n + K* – 2. If 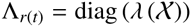 is a diagonal rate matrix relating the ordered parameter spaces to the coalescent rates with *j*^th^ component Λ_*r*_(*t*)[*j*], then the Snyder filter can be described as:

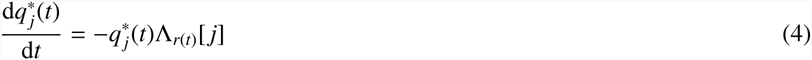

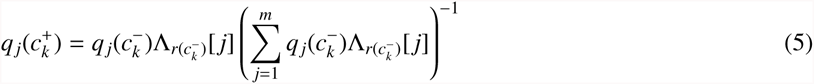

Equation 4 describes the evolution of the posterior between event times and admits a matrix exponential solution on the probability components. At event times this solution is discontinuously renormalised by the rate matrix, as in equation 5, with 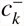 and 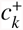 as infinitesimally before and after the *k*^th^ coalescent event, such that 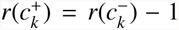. When *K* > 1 (heterochronous sampling) an extra condition needs to be included to account for *r*(*t*) falling to 1 after some time *τ*, where *τ < s*_*K*_ (the last sample time). This condition can be written as: {*q*(*τ < t < s*_*i*_): [*r*(*t*) = 1] Λ [*i* ≤ *K*: *s_i_> t*]} = *q*(*τ*). This means that the posterior, at *τ*, is maintained until the next sampling time, at which point *r*(*t*) rises above 1 again. This follows because over this time period the rate Λ_*r*(*t*)_ [*j*] = 0 so that equation 4 gives 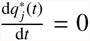 for 1 ≤ *j ≤ m*.

As mentioned previously, if equation 4 is written in terms of normalised probabilities, it becomes non-linear. This version is: 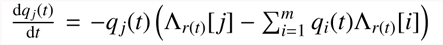 [17] with the sum contributing the non-linearity. If we define *φ*_*j*_ = –Λ_*r*_(_*t*_)[*j*], with the average ⟨*φ*⟩ taken across the current posterior *q*(*t*), then we can write 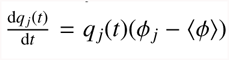. The Snyder equation therefore describes movement away from a local mean with time.^3^

The conditional estimates of 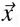 and 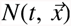 can be computed from the posterior, *q*(*t*. The sum in equation 6 indicates marginalisation across *l* – 1 dimensions and *A*^T^ denotes the transpose of *A*. The conditional mean estimate, 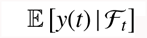, minimises the mean square error (MSE) index, *R*_*t*_, of equation 8 and uses all the causally available information, 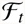. Here *y*(*t*) is some function of 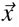 such as *x*_*i*_(*t*) or 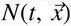 and *y*_*e*_(*t*) is some arbitrary estimator of *y*(*t*).

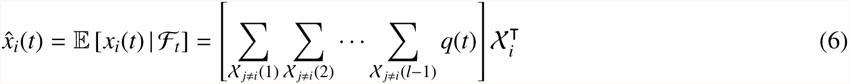

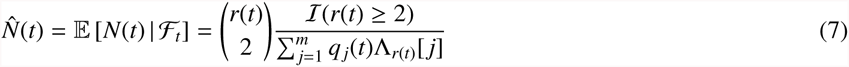

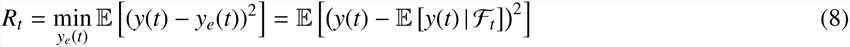

Thus the filter takes a time series of sampling and coalescent events, 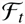, and parameter priors, 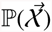, as its input, and iterates to a posterior 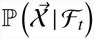 from which conditional mean estimators are obtained. Estimating *N*(*t*) is equivalent to estimating *λ*(*t*) since *r*(*t*) is known from 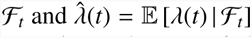. Since the coalescent model has a fixed parametric form in this framework, the optimal estimate that makes use of all the information exists at the end of the event time series. This is at *t = T = c*_2_, which is the time of the most recent common ancestor (TMRCA) of the samples. Thus, substituting *q*(*T*) for *q*(*t*) in the above equations gives the minimum mean squared error (MMSE) estimators conditioned on 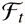. This final posterior is used in the results section below, and has MMSE *of R*_*T*_.

The filter mechanics remain largely unchanged even when generalised to demographic functions that are much more complex than those examined here. For example, if *N*(*t*) is stochastic and described by a continuous time Markov process that has its states as the parameters to be inferred, then it is only necessary to replace *Λ*_*r*_(_*t*_) with Λ*r*_(*t*)_ **–** *Q* in equation 4. *Q* is the infinitesimal generator of the Markov process. The filter can also be simplified to perform inference for normal inhomogeneous processes, or combined into a more involved form that can handle complex Poisson processes with feedback dependent and stochastic multimodal rates [22]. These generalisations would, in theory, allow the filter to incorporate extra demographic information from other time series sources. Future research will explore these possibilities.

## 3. Results

We apply Snyder filtering theory to several coalescent inference problems. We start by examining the Kingman coalescent, for which we derive an analytical MMSE estimator. We also show the relation between Snyder estimation and ML inference. Next, we simulate genealogies under different demographic functions and use these models to explore the performance and flexibility of Snyder based inference under both isochronous and heterochronous sampling. Lastly, we adapt the filter to a well studied dataset of Egyptian hepatitis C viral sequences and compare our results with those of previous analyses [7]. In the Appendix we complete our analysis by developing the connection between the Snyder filter and the classic skyline plot.

### 3.1. The Standard Kingman Coalescent has an Analytic Solution

The standard Kingman coalescent has a constant *N*(*t*) = *x*_1_ and is isochronously sampled, hence *r*(*c*_*k*_) = *k*. If *δ*_*k*_ = *c*_*k*-1_ **–** *c*_*k*_ is the waiting time for the Kingman process to transition from *k* to *k **–*** 1 lineages, then *δ*_*k*_ follows an exponential distribution with rate 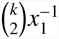. We show that, for the Kingman coalescent, we can reformulate the data structure of this problem to allow a direct analytical treatment of the Snyder filter that does not require solving equations 4 and 5. This reformulation (and analytic treatment) does not work for more complex coalescent models.

Let 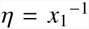 and consider a single coalescent tree with *n* tips and *n* – 1 coalescent waiting times of duration *δ*_*k*_, 2 ≤ *k* ≤ *n*. As in [29], the likelihood function, *L*_1_(*η*), follows due to the independence of the coalescent intervals. The sufficient statistic 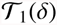 can then be Fisher-Neyman factorised (into a product of functions *h*_1_ and *g*_1_) and the ML estimator, 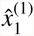, derived. Note that 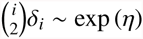. While in general 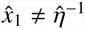. it is valid here because the same value of 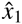 maximises the partial derivatives of the likelihood with respect to *η* or *x*_1_.

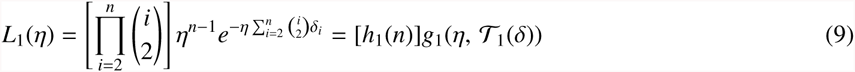

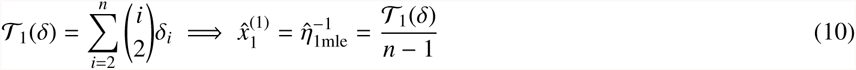

Now, let us assume a single *j*^th^ coalescent event is observed from each of *n –* 1 independent trees (unlinked loci). The data are now *τ*_*j*_(*i*) for 2 ≤ *i ≤ n* (*i* starts from 2 for notational consistency). By the properties of exponential scaling, the product 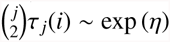. The likelihood, *L*_2_(*η*), sufficient statistic, 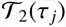 and ML estimator 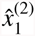 follow similarly.

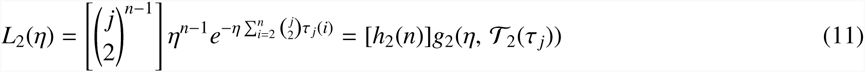

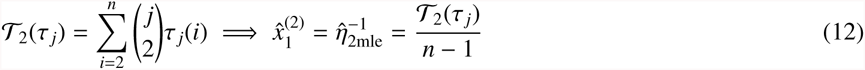

Since both sufficient statistics, 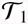 and 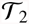, comprise a sum of n – 1 independent exponential variables with rate *η*, and the corresponding ML estimators only depend on the sufficient statistic and the sample size, *n*, then it is equally efficient to sample coalescent waiting times from either a single tree or from multiple trees. Further, 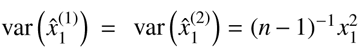 [29]. Hence, for a constant sized population, the *n* – 1 coalescent times of one tree contain the same information as the *j*^th^ coalescent times from *n –* 1 independent trees (unlinked loci).

As a result, the standard Kingman coalescent can be reformulated from a self-correcting to a simple homogeneous Poisson process. In the context of the Snyder filter (equations 4–5), this results in the rate matrix *Λ*_*r*(*t*)_ being replaced by the constant matrix Λ = diag (*λ* (*X*_1_)). *X*_1_ is the space within which the true value *x*_1_ lies. This transformation means that the explicit result derived in [17] for the posterior probability, 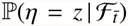 under a constant intensity Poisson process with prior ℙ(*η* = *z*), is applicable to the Kingman coalescent (equation 13), observed until *t* into the past. We use the 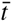 notation to indicate the scaled time which corresponds to the actual time *t* into the past (the scaling is by 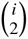 for periods of *i* existing lineages). Here *z* is an arbitrary variable that spans the parameter space and 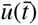 is a coalescent event counting function on scaled time. By the information equivalence previously mentioned in equations 10 and 12, 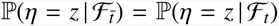 and 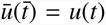. The general analytic MMSE estimator, 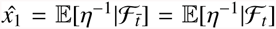 is then given by equation 14. If we assume a uniform prior on *x*_1_, then we can obtain a closed form solution for the MMSE estimator, as in equation 15, by recognising the Laplace transform definition: 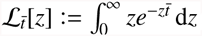.

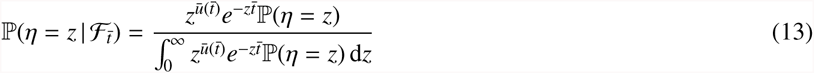

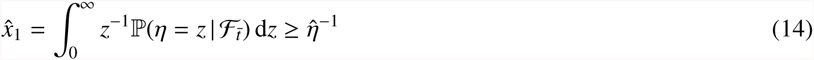

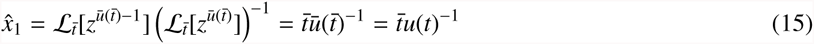

As far as we are aware, these are the first explicit MMSE solutions for Bayesian inference on the Kingman coalescent. These expressions depend on a scaled time which is a transform of the actual observed coalescent stream length. Applying Taylor’s expansion and noting that any observed set of Kingman coalescent event times is equally likely for a sufficiently large *n* (hence the second equality in equation 17), allows an approximate expression for the MMSE, *R*_*t*_, to be derived (in normal time).

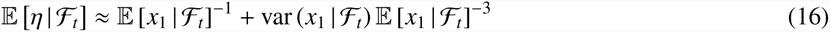

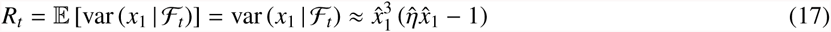

In the limit of *t →* ∞ (an infinitely long observed coalescent information stream 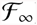), *x*_1_ → *η*^−1^ so that *R*_∞_ = 0, as expected. Similar analytic expressions are not available when the demographic function varies with time because it is not possible to scale the exponential parameters so as to remove the count-dependent component of the process.

Expression 13 allows us to also derive an informative link between the Kingman coalescent MMSE solution and standard ML analysis (which would involve maximising the single tree likelihood of equation 9 to yield the estimator of equation 10). Consider the final Snyder posterior at time, *T*, when the last coalescent event has occurred (the TMRCA of the sample). This corresponds to scaled time 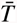 and satisfies 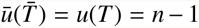. Equation 13, with some constant *F*, then becomes equation 18. If we set 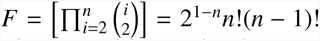, and note that 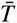 is the same as the sufficient statistic 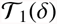, then we can apply Bayes theorem with equation 9 to get equation 19.

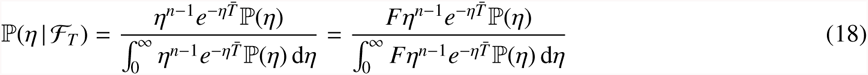

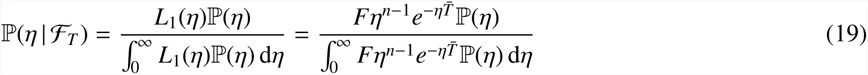

If a uniform prior is assumed then we can also substitute into equation 15 and recover 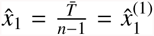 from equation 10. Thus Snyder and standard ML inference give the same estimation result for the Kingman coalescent when all coalescent events up to the TMRCA are evaluated.

### 3.2. Snyder MMSE Estimation of Simulated Demographic Models

We now explore the performance of the complete Snyder filter (described by equations 4-5) by applying it to data simulated under several demographic models that have been used in the epidemiological literature. A demographic model with *l* ≥ 1 parameters, is once again denoted 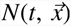 (or just *N*(*t*) for short) with *x*_*i*_ being a true parameter value that must be inferred from a discrete search domain of *m*_*i*_ values, *X*_*i*_. Coalescent event times from a simulated tree with *n* tips are generated using standard approaches such as time rescaling [30] or rejection sampling. The resulting *n–*1 coalescent event times for each tree are then iteratively processed by the Snyder filter, resulting in the sequentual calculation of the final joint posterior, *q*(*T*). The prior and posterior are defined over an *m* valued space, 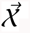 (see section 2.2). MMSE conditional estimates of the parameters 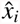 and of the overall demographic history 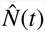 are obtained by either marginalising *q*(*T*) or an appropriate function of it.

Three main demographic functions are simulated and analysed here. The first is exponential growth *N*(*t*) = *x*_1_*e*^−x2*t*^ [31], which describes a rapidly rising population (in forward time). Here *x*_1_ = *N*(0)is the population size at the present and *x*_*2*_ is the exponential growth rate. We used a time rescaling algorithm to iteratively generate the *k*^th^ coalescent time, *c*_*k*_, from the (*k* + 1)^th^ via: 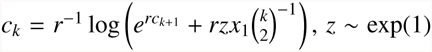. The second demographic function is the logistic growth model which represents density dependent population growth and is defined as 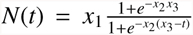 [12]. An offset parameter, *x*_4_, is added so that *N(t) ≥* 0 for all *t >* 0. Parameters *x*_1_ and *x*_2_ are defined as in the exponential growth model and *x*_3_ is the half life. The logistic model was simulated using a rejection sampling algorithm that generates points from a homogeneous process with rate 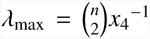 and then chooses one as the next coalescent event with acceptance probability 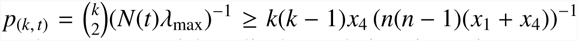. Lastly, we used a sinusoidal function, *N*(*t*) = *x*_1_ sin(x_2_*t* + *x*_3_) + *x*_4_, to model cyclical population dynamics. Here *x*_1_ is the cycle amplitude and *x*_2_ its frequency of occurrence, with phase and magnitude offsets given by *x*_3_ and *x*_4_ respectively. Data from this model was also generated using a rejection sampling algorithm with 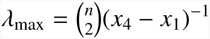 and 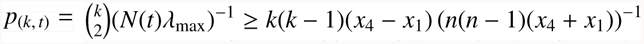.

For every demographic model, we simulated a total of *M* = 1000 trees, each with *n* = 200 tips and then applied the Snyder filter to the coalescent times of each tree to obtain informed joint posterior distributions. We set the true values of each parameter at approximately the midpoint of its space so that it would be described by the prior. Defining *T*_*i*_ as the final observation time for the *i*^th^ tree (in this problem we will always observe until the TMRCA) then the Snyder filter yielded 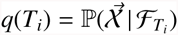 (see section 2.2), from which the square error between the true and estimated value of some quantity of interest, *y(T*_*i*_) was calculated. Here *y(T*_*i*_) was either the demographic function or its parameters. We averaged these values over the *M* trees to obtain the following frequentist approximation to the MMSE of equation 8, *R*_*r*_(*y*), and the relative percent MMSE, *J*_*r*_(*y*).

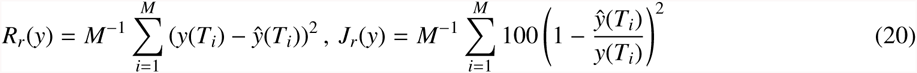

The distributions of the relative square errors of estimated demographic parameters for the exponential, logistic and sinusoidal models (with mean *J*_*r*_) are given in Figures 1c–d, 2a–d and 3a–d respectively. Figures 1b, 2e and 3e provide an illustration of the Snyder reconstruction of the demographic function for each model, with estimation bounds delimited by twice the standard deviation of the posterior. For the exponential model we also show the bivariate joint distribution of its parameters in Figure 1a.

**Figure 1.**
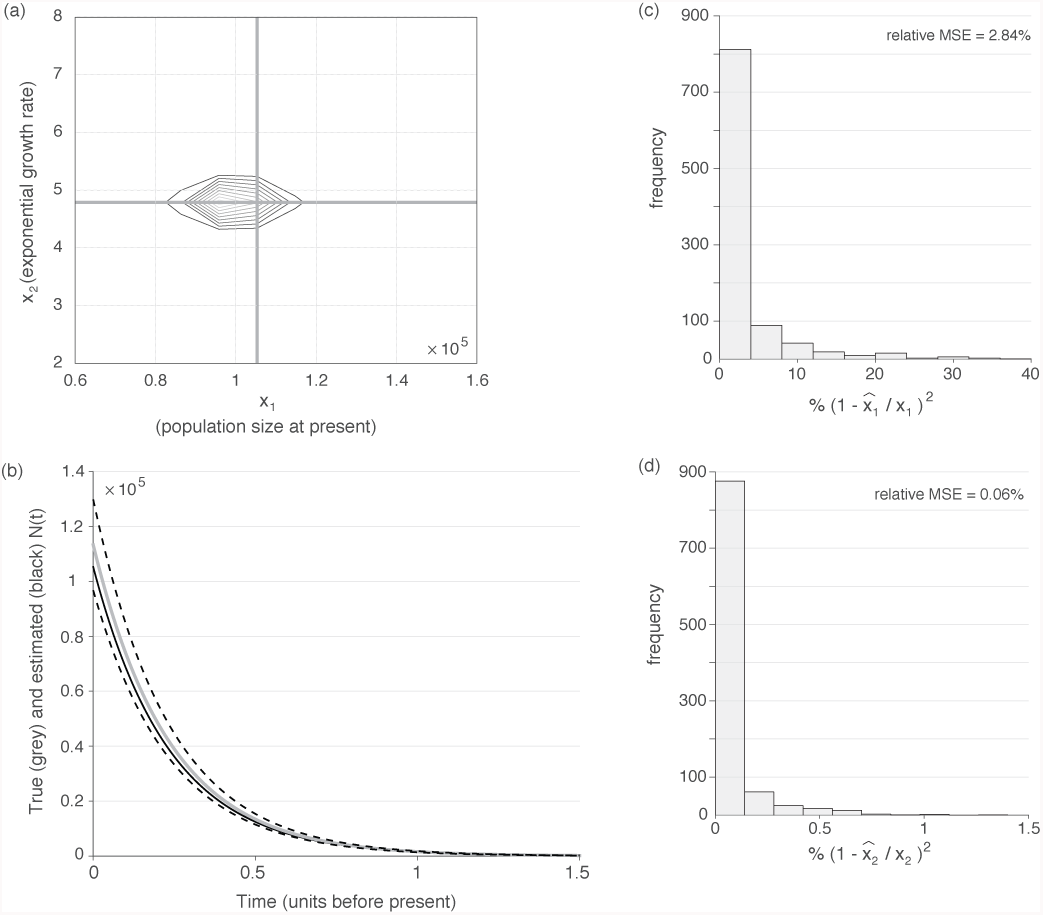
Snyder estimates under the exponential growth coalescent model. *N*(*t*) = x_1_*e*^−*x*2*t*^. a) Contour plot of the joint posterior for the two model parameters, *x*_1_ and *x*_2_. Contour lines show values of the posterior P(*x*_1_, *x*_2_ | data). Thick grey lines show the true values of the two parameters. (b) A reconstruction of the estimated demographic function, obtained from a single tree with 200 tips simulated under exponential growth. The true demographic function is in grey and the estimated function is in black. Dotted lines are uncertainty bounds derived from twice the standard deviation of the posterior. (c)-(d) Histograms of the relative square estimation errors of each model parameter, measured across 1000 replicate trees. Simulations were done at: [*m*_*i*_, *m*, *n*, M] = [20, 400, 200, 1000].

**Figure 2.**
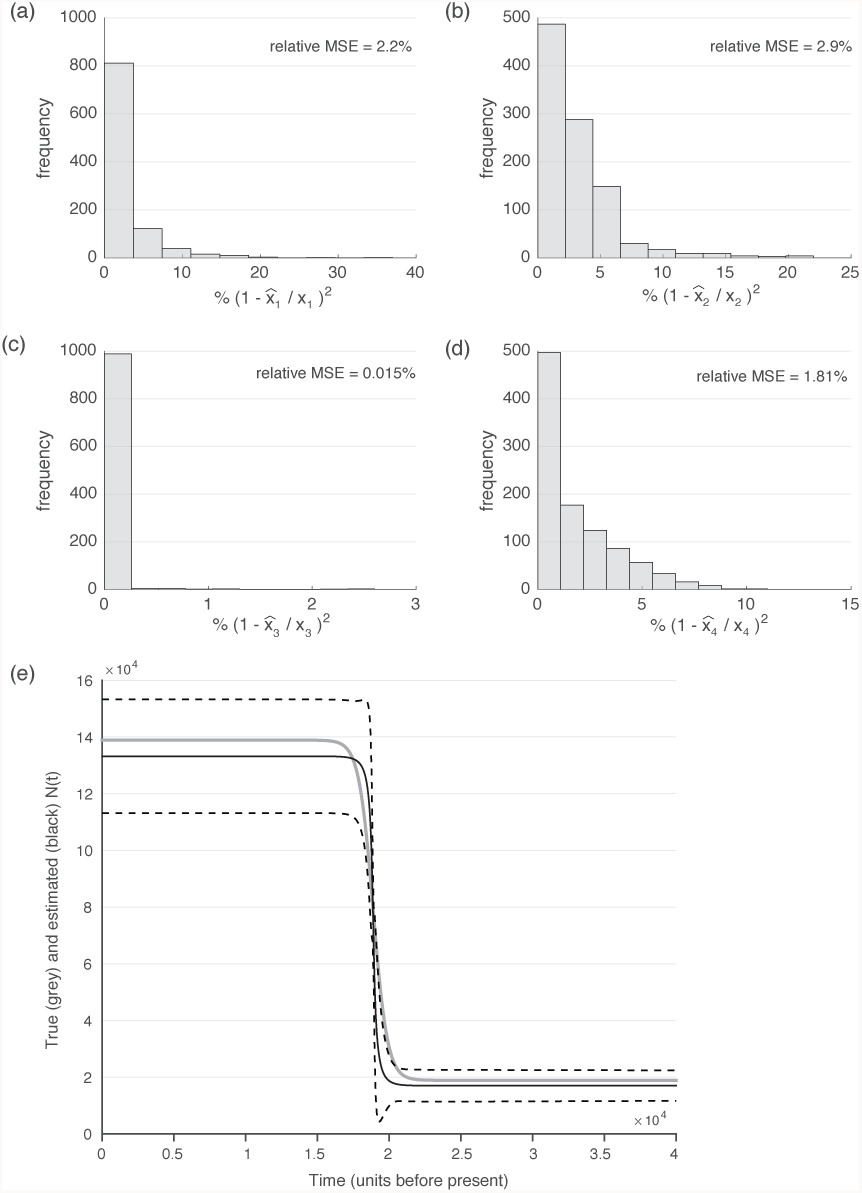
Snyder estimates under the logistic growth coalescent model. 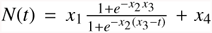. (a-d) Histograms of the relative square estimation errors of each model parameter, measured across 1000 replicate trees. (e) A reconstruction of the estimated demographic function, obtained from a single genealogy simulated under rapid logistic growth. The true demographic function is in grey and the estimated function is in black. Dotted lines are uncertainty bounds derived from twice the standard deviation of the posterior. Simulations were done at: [*m*_*i*_, *m*, *n*, *M*] = [10, 10^4^, 200, 1000]. An *m*_*i*_ of 20 was used in (e).

**Figure 3.**
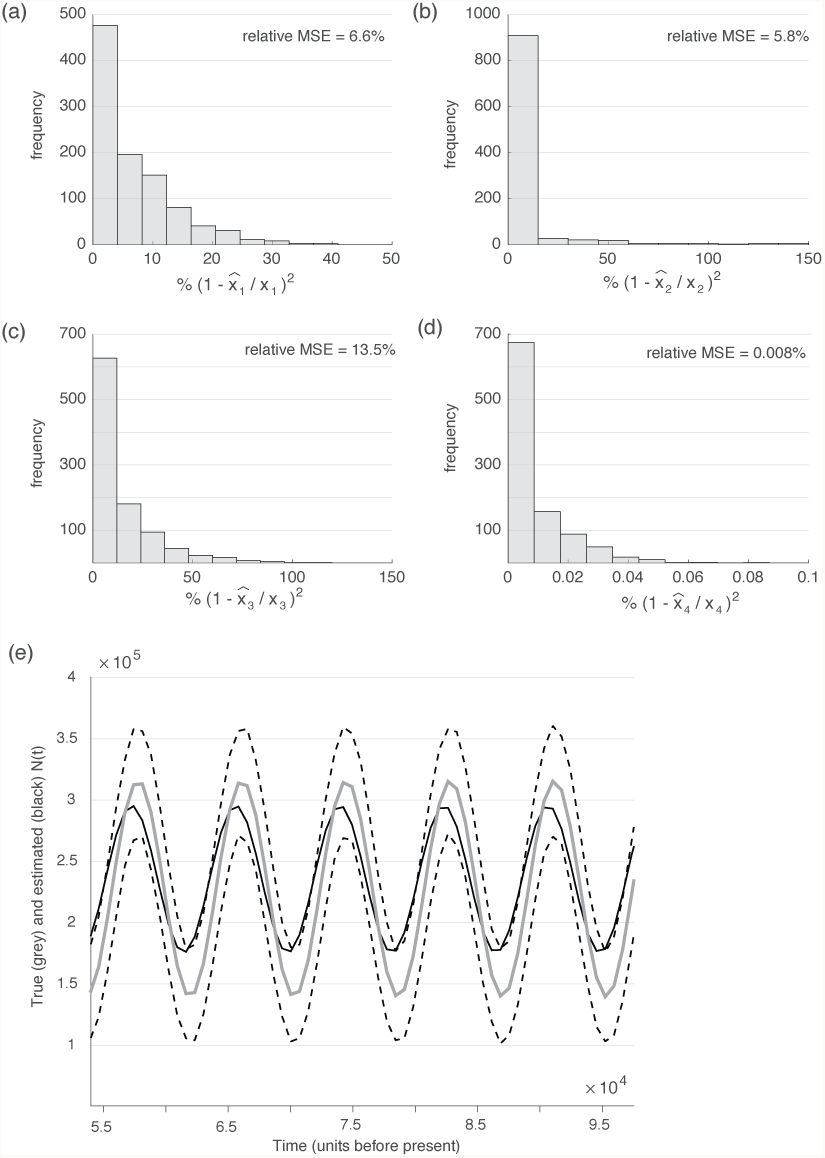
Snyder estimates under a sinusoidal coalescent model. *N*(*t*) = *x*_1_ sin (*x*_2_*t + x*_3_) +*x*_4_. (a-d) Histograms of the relative square estimation errors of each model parameter, measured across 1000 replicate trees. (e) A reconstruction of the estimated demographic function, obtained from a single genealogy simulated under the sinusoidal model. The true demographic function is in grey and the estimated curve is in black. Dotted lines are uncertainty bounds derived from twice the standard deviation of the posterior. Simulations were done at: [*m*_*i*_, *m, n, M*] = [10, 10^4^, 200, 1000].

### 3.3. Extension to Demographic Models with Heterochronous Sampling

The preceding simulations considered only isochronous sampling in which all the sequences were sampled only at the present (*t* = 0). Here we show how Snyder filtering can be applied to the heterochronous coalescent process. We examine two demographic functions that involve rapid epidemic growth forward in time. The exponential growth model [31]: *N*_1_(*t*) = *x*_1_*e*^*−x*2*t*^, as introduced in section 3.2, and the constant-exponential-constant (con-exp-con) model [7]: 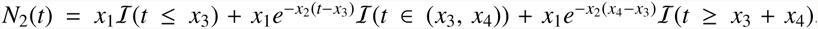. *N*_2_(*t*) is simply a translated and time limited version of *N*_1_(*t*) so that 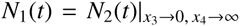. The con-exp-con function can be considered as a piecewise equivalent of the logistic function from section 3.2 with parameters x3 and x4 controlling the times at which the population switches between phases of constant size and exponential growth.

We simulate the coalescent event times by modifying the time rescaling technique in [30] to allow for the extra conditions that serial sampling of the tips impose. The key step of this algorithm involves solving for *c*_next_, the possible next coalescent time. If *c*_next_ is less than the next sample time then the lineage count is reduced by 1 since the next event is a coalescent one. If it is larger, then the time is set to the subsequent sampling time, the number of lineages updated with the new sample count, and the process of calculating *c*_next_ restarted. Since the simulated process is stochastic, the number of lineages could fall to 1 before the last sample time. In this case, the coalescent rate is 0 until the next sample time and the posterior distribution stays unchanged over this interval (see section 2.2 for the mathematical expression of this condition). The specific time rescaling solution for the con-exp-con model is given below with *z* ~ exp(1). The exponential demographic function only requires the solving of the last condition with *x*_3_ = 0 and *x*_4_ → ∞.

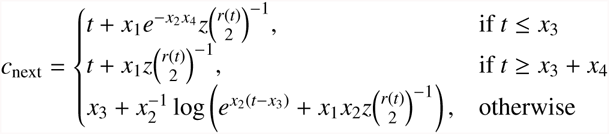

Having generated the simulated data 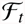, we then used the Snyder filter to compute the posterior, 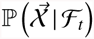. We simulated *N*_1_(*t*) and *N*_2_(*t*) over identical 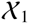 and 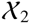 spaces using *n* sampled lineages introduced uniformly at times *s_i_ = (i –* 1)T_samp_*K*^−1^, 1 ≤ *i* ≤ *K*, for some maximum sampling time *T*_samp_. At each sample time *n*_*i*_ = *nK*^−1^ = *n*\* lineages were added. Figure 4a illustrates Snyder filter estimation from heterochronous trees with *n* = 200 tips, sampled at *K* = 10 times, for both the exponential growth and con-exp-con demographic models. For the latter, *T*_samp_ = *x*_3_ + (*x*_3_ + *x*_4_) is used so that sampling times are symmetric with respect to the period of exponential growth. The Snyder filter estimates both functions well. Figure 4b shows that the exponential growth rate and initial population parameters are better estimated for the exponential model than for the the con-exp-con one. This is expected, because *N*_2_(*t*) has a much larger parameter search space than *N*_1_(_t_).

**Figure 4.**
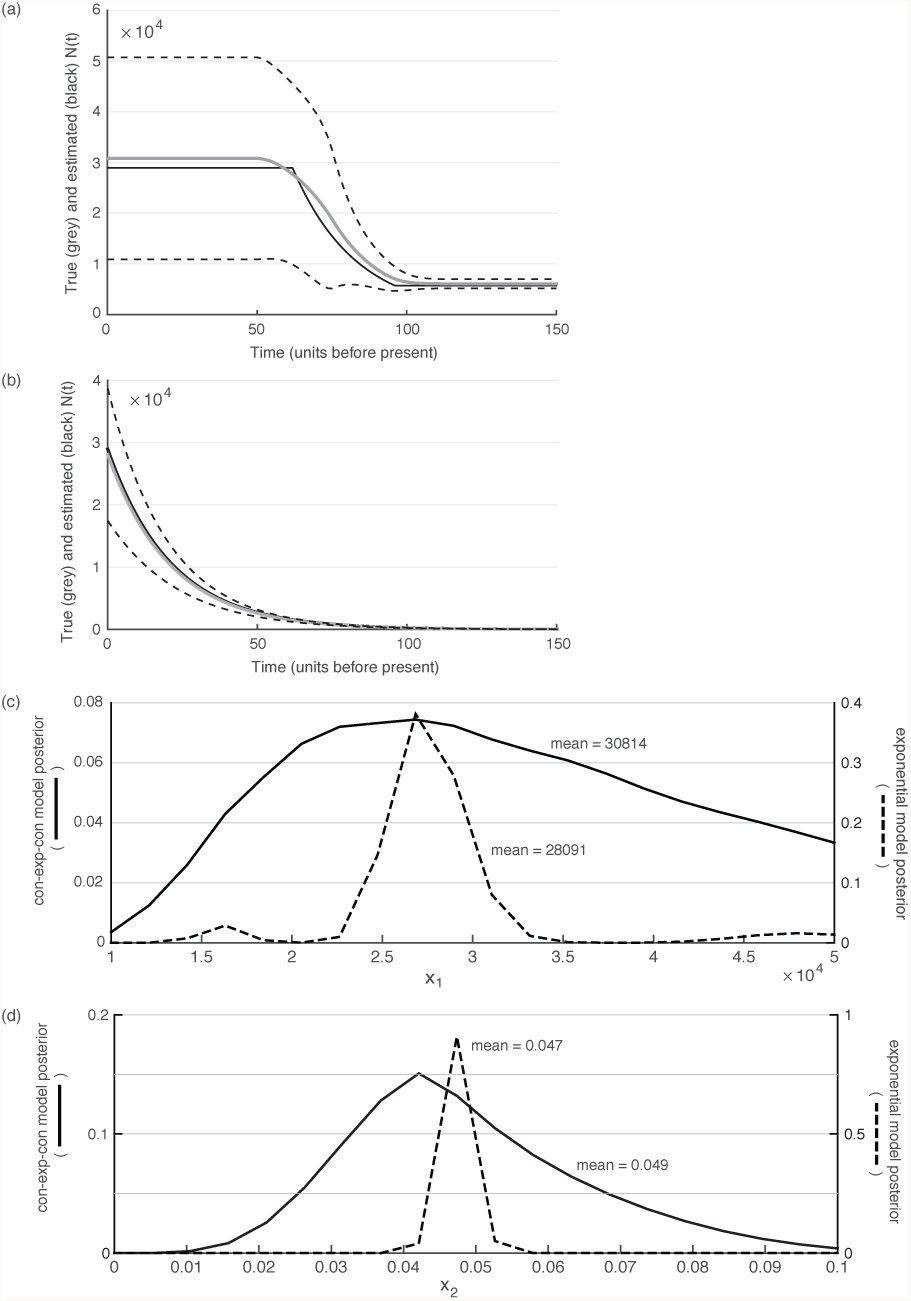
Snyder estimates under heterochronously sampled exponential and con-exp-con models. (a)-(b) Reconstruction of the con-exp-con and the exponential demographic functions, respectively. In both cases the simulated populations was heterochronously sampled 10 times, uniformly across the time range. Each sample introduces *n**\**** = 20 lineages so that *n* = 200. The true demographic function is in grey and the estimated curve is in black. Dotted lines are uncertainty bounds derived from twice the standard deviation of the posterior for each model. (c)-(d) Posterior distributions for the initial population (*x*_1_) and exponential growth (*x*_2_) parameters, respectively. The con-exp-con model (solid line) has a much larger parameter space than the exponential model (dotted line) which explains the reduced certainty and wider posterior in the estimates of the former. Simulations were done at: [*m*_*i*_, *n, K*] = [20, 200, 10].

### 3.4. Estimation of the Hepatitis C (HCV) epidemic in Egypt

Having tested the Snyder filter performance on simulated data, we now illustrate its application to empirical data. One particular data set, comprising hepatitis C virus (HCV) gene sequences sampled in Egypt, has been repeatedly used as a benchmark for the performance of parametric and non-parametric coalescent estimators in molecular epidemiology (see [15] [32] [13] [33] [34]). It therefore enables different inference approaches to be directly compared. A further benefit of this data set is that its demographic history is partially known through independent epidemiological information. Specifically, the very high current prevalence of HCV in Egypt is very likely due to the rapid transmission and amplification of this blood-borne virus during a widespread public health campaign of parenteral antischistosomal therapy (PAT). Poor sterilisation of needles used to deliver the drug treatment led to the inadvertent transmission of HCV [36]. These injections were primarily given between 1920 and 1980. Consequently, we would expect coalescent inference methods to reconstruct a rapid rise in the effective population size of HCV during this period. The Egyptian HCV dataset comprises 63 HCV genotype 4 gene sequences, 411 nucleotides in length, sampled isochronously in 1993 [37]. In previous work [7] a con-exp-con model was fitted to this dataset using a Bayesian MCMC approach. In order to achieve comparative results, we apply the Snyder filter to exactly the same demographic function. The con-exp-con demographic model is the same as that introduced in section 3.3, but with the indicator functions expanded. Setting *x*_1_ = *N*_*C*_, *x*_2_ = *r*, *x*_3_ = *x* and *x*_4_ = *y* – *x*, this demographic model can be written as in [7] with *t* > 0 describing time in the past from 1993 (the time of sampling).

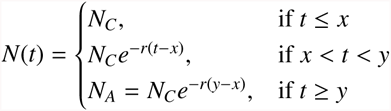

To obtain coalescent times suitable for Snyder filter inference, we estimated a molecular clock phylogeny from the Egypt HCV sequences as follows. First, we used the software Garli [38] to estimate a maximum likelihood phylogeny for the sequences under the general time reversible nucleotide substitution model with gamma distributed site rate heterogeneity. This was the model previously used in [7]. Next, we used the program R8s [39] to convert the ML tree into an ultrametric phylogeny whose branches are scaled in units of years. This conversion was performed using a Langley-Fitch 3-rate molecular clock model, and the evolutionary time-scale was enforced by constraining the TMRCA of the tree to the range of values reported in [7]. The resulting time-scaled tree is shown in Figure 5a. We extracted the desired coalescent times from this tree and then applied the Snyder filter with [*m*_*i*_, *m*] = [20, 20^4^]. Prior distributions were set to match those of [7] as closely as possible. The resulting marginalised posteriors are given in Figure 5b and are compared to the posteriors obtained by Bayesian MCMC sampling in [7]. The Snyder MMSE estimate of the demographic function is shown in Figure 5c. It is clear that the Snyder filter gives results which are consistent with the known epidemic history of HCV in Egypt and similar to those of previous studies [7].

**Figure 5.**
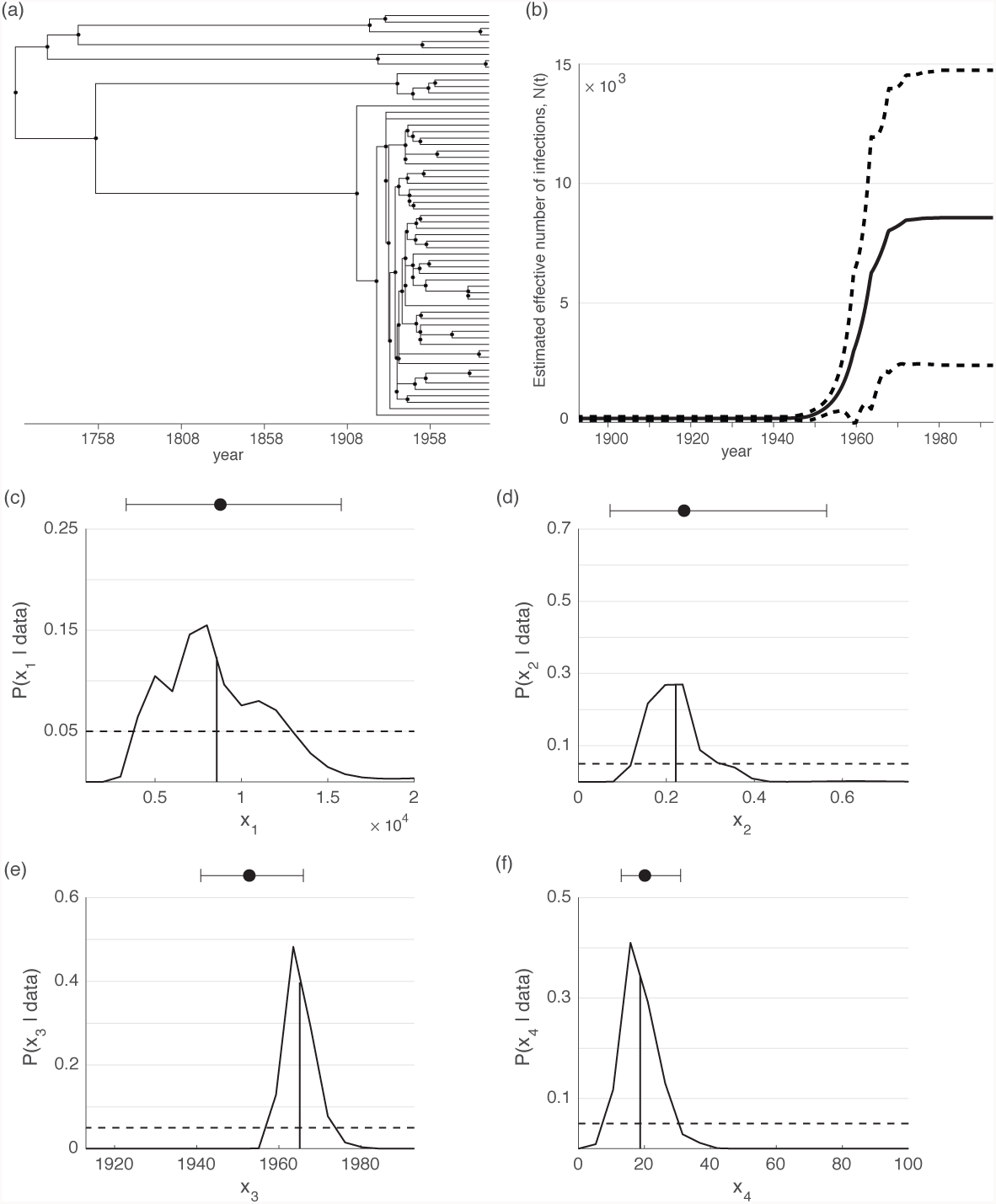
Snyder estimates of the HCV epidemic showing exponential growth in the early 20^th^ century. (a) The ultrametric time-scaled tree derived from the 63 sequence HCV dataset using Garli and R8s. The branching times of this tree provide the coalescent event times for Snyder filtering. (b) The Snyder estimate of HCV population size using a con-exp-con demographic model. The continuous black line is the conditional mean estimate while the dotted lines are delimited by twice the standard deviation of its posterior. Rapid growth during the PAT period (1920–1980) is clear. (c)-(f) Marginal posteriors for each parameter of the con-exp-con demographic model (solid black line). The dotted lines are the uniform priors used. The vertical black lines are the conditional mean estimates for each parameter. The Snyder estimates and their uncertainty compare well with those reported in [7]. The parameter estimates from [7] are based on a Bayesian MCMC sampling technique and are given by the boxplots above each posterior. The Snyder filter used: [*m*_*i*_, *n*] = [20, 63].

## 4. Discussion and Conclusion

In this paper we introduced the point process Bayesian Snyder filter as a viable technique for coalescent inference and we evaluated its performance on several demographic models. By reinterpreting the variable population size coalescent process under heterochronous sampling as a self-correcting Poisson process, we showed that it was possible to apply the Snyder filter to coalescent estimation problems. Further, for the standard Kingman coalescent, we were able to derive an analytic solution for the Synder MMSE estimator by recognising the equivalence between coalescent data from a single tree and multiple loci. This solution was equivalent to a maximum likelihood estimate and clarified the relation between Bayesian and ML approaches.

We then applied the Snyder filter to isochronous data simulated under three different parametric demographic functions. Two of these functions (exponential and logistic) are commonly used to model viral population growth whilst the third (sinusoidal model) is consistent with the type of cyclical epidemic behaviour exhibited by influenza and other viruses. In all cases, the Snyder filter correctly inferred the underlying dynamics and achieved good MMSE estimates of the model parameters. We then showed how Snyder filtering could be readily extended to heterochronous sampling and compared estimates across related exponential and con-exp-con demographic functions for illustration. Once again estimates were accurate and the impact of a larger parameter space was observed by comparing estimate uncertainty between these models. A key benefit of the Snyder filter is its flexibility and ease of use. This was apparent for all of these simulations, as the filter algorithm remained unchanged across all the isochronously and heterochronously sampled models, beyond specification of the demographic function and its priors.

We then assessed filter performance on empirical data from the Egyptian HCV epidemic. In keeping with previous studies on this dataset, we used a parametric con-exp-con demographic model. The filter achieved estimates that compared well with those previously obtained using Bayesian MCMC sampling [7]. It is worth noting that the Bayesian MCMC sampling approach incorporated uncertainty in the phylogeny and the molecular clock model, whereas we assumed that the coalescent event times were known without error. Further work will be necessary to understand how these sources of error can be best incorporated into the Snyder filter framework.

Thus the Snyder filter has the potential to be a capable, alternative inference method for coalescent inference. Since it only involves the solution of linear differential equations, it is easy to implement and provides stable estimates (the filter is generally not susceptible to numerical explosion or lack of convergence). Further, because it allows straightforward calculation of conditional means, it easily leads to optimal (MMSE) estimates. Hence the Snyder filter may serve as a useful benchmark for testing the limits of other estimation schemes, and for providing bounds on achievable estimation accuracy [23].

However, we note that while its implementation is linear, the Snyder filter is a non-linear MMSE filter (its linear equations must be normalised). The importance of this is apparent when the parametric Snyder filter is compared to the non-parametric classic skyline plot. In the Appendix, we demonstrate that the classic skyline plot is a minimal information linear estimator for the coalescent process. This can be improved to a linear MMSE estimator if more information is introduced via first and second order process statistics. If this linearity constraint is maintained, no further improvements (defined as reductions in estimation MSE) are possible, even if more coalescent process information is available [24]. At this point the MSE can only be further minimised by allowing for non-linear filter formulations that make use of extra information, usually embedded in the form of some demographic model structure (parameters). The parametric Snyder filter is exactly the non-linear MMSE estimator that makes maximum use of this additional coalescent process information. These connections hint at the reasons underlying the trade-off between estimation accuracy and methodological simplicity that was mentioned in the Introduction.

The filter was originally developed for doubly stochastic Poisson processes [24]. As a result, it should be capable of handling stochastic demographic functions, or input streams with mutiple types of stochastic events (as, for example, might be encourtered within the structured coalescent model). The Snyder filer could also be extended to use alternate descriptors of the relation between population dynamics and phylogenetic structure, such as birth-death models [40]. Extending the filter to these cases will form the basis of future research. The flexibility, exactness and robustness of the Snyder filter suggest this approach has much promise. We expect that its benefits will only become fully apparent when it is applied to more complex models that are computationally challenging to implement in other frameworks, such as Bayesian MCMC sampling.

## Acknowledgements

This work was supported by the European Research Council under the European Commission Seventh Framework Programme (FP7/2007-2013)/European Research Council grant agreement 614725-PATHPHYLODYN.

## Appendix The Relation between the Snyder Filter and the Classical Skyline

Consider the coalescent process with deterministically time varying population size. In this appendix, we focus on characterising the relationship between the Snyder filter and non-parametric coalescent estimation schemes for this process. The classic skyline plot, developed in [12] is a well known non-parametric method for estimating coalescent demographic functions. For a given coalescent waiting time *δ*_*k*_ = *c*_*k–1*_ – *c*_*k*_, starting with *k* lineages, the skyline estimate, 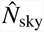 is given below, with 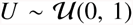 and *W = –* log (*U*) ~ exp(1) ≡ Gam(1, 1). It is an estimate of the harmonic mean of the population, 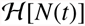, over each event interval, corrupted by multiplicative noise, *W*.

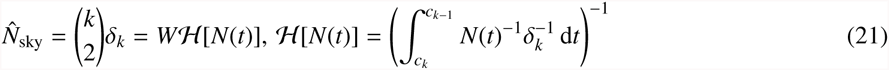

Define *η*(*t*) = *N*(*t*)^−1^ as the intensity to be estimated and let 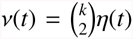. Note that 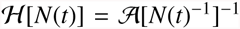 where 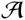 is a functional that calculates the arithmetic mean. The skyline then implies an estimator 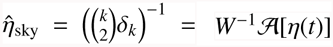 with the multiplicative noise now being *W*^Ȓ1^ ~ Inverse Gam(1, 1). In [24] an expression is given for linear filters of doubly stochastic Poisson processes. When scaled with 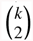, all linear filters of *η*(*t*) over *δ*_k_ satisfy equation 22 for arbitrary functions *g*(*t*) and *h*(*c_k_, τ*). Here *u*(*τ*) is the usual counting process for coalescent events up to time *τ* ≤ *t* and 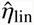 is the intensity estimate resulting from the linear filter.

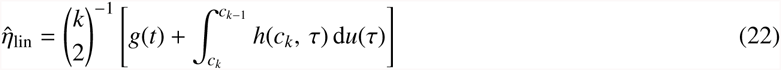

Choosing *g*(*t*) = 0 and 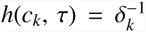 results in 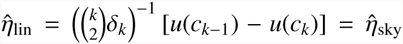. The classic skyline plot is therefore a type of linear filter for doubly stochastic Poisson processes. More specifically, it is a moving average filter that only requires knowledge of the interval time between coalescent events, which serves as the averaging time. Generalisations of the skyline method essentially alter the construction of the integral in equation 22 [6].

The benefit of moving average filters is that they require minimal process information. If the first and second order statistics of the intensity process are also known then a linear MMSE estimator can be obtained which outperforms all possible other linear filters. Provided the estimator is constrained to be linear this MMSE filter cannot be improved upon, even with extra knowledge of higher order statistics or structural information (such as a parametric model). The linear MMSE estimator is equivalent to the optimal estimate of an intensity, given that it was corrupted by additive Gaussian noise [24]. The Snyder filter generalises the linear MMSE filter for doubly stochastic Poisson processes by removing the linearity constraint. This allows extra information from a parametric model to be used and leads to superior performance (smaller MSE). In summary, the skyline is non-parametric, linear and makes use of a minimum of coalescent information. Given mean and covariance information of the coalescent intensity, the skyline can be adapted to a linearly constrained MMSE filter. If a coalescent rate model is known then this can be further improved with the (unconstrained) Snyder non-linear MMSE estimator. Beyond this, no performance improvements are possible without accounting for genealogical uncertainty.

## Footnotes

3 This form contains an interesting conceptual detail. If *φ* can be considered a fitness vector then this is a continuous time replicator equation from evolutionary game theory [26]. Further, the discontinuous update in expression 5 obeys the discrete time version of the same replicator equation as step size, away from an event time, becomes infinitesimally small. The correspondence between Bayesian statistics and the replicator equation has been identified in discrete time [27] [28]. Here we note, for the first time (to our knowledge), that the Snyder filter presents a continuous time analogue. The Snyder coalescent solution may therefore be seen as the evolution of a series of *m* strategies with fitness given by the negative of the coalescent rate (which acts like some measure of stability).

